# Enveloped viruses pseudotyped with mammalian myogenic cell fusogens target skeletal muscle for gene delivery

**DOI:** 10.1101/2023.03.17.533157

**Authors:** Sajedah M. Hindi, Michael J. Petrany, Elena Greenfeld, Leah C. Focke, Alyssa A.W. Cramer, Michael A. Whitt, Vikram Prasad, Jeffrey S. Chamberlain, Benjamin Podbilewicz, Douglas P. Millay

## Abstract

Entry of enveloped viruses into cells is mediated by fusogenic proteins that form a complex between membranes to drive rearrangements needed for fusion. Skeletal muscle development also requires membrane fusion events between progenitor cells to form multinucleated myofibers. Myomaker and Myomerger are muscle-specific cell fusogens, but do not structurally or functionally resemble classical viral fusogens. We asked if the muscle fusogens could functionally substitute for viral fusogens, despite their structural distinctiveness, and fuse viruses to cells. We report that engineering of Myomaker and Myomerger on the membrane of enveloped viruses leads to specific transduction of skeletal muscle. We also demonstrate that locally and systemically injected virions pseudotyped with the muscle fusogens can deliver micro-Dystrophin (μDys) to skeletal muscle of a mouse model of Duchenne muscular dystrophy. Through harnessing the intrinsic properties of myogenic membranes, we establish a platform for delivery of therapeutic material to skeletal muscle.

## Introduction

Membrane fusion is critical for the movement of molecules between cellular compartments, fertilization, development of multinucleated cells, and viral infection (Martens and McMahon, 2008). The fusion of two membranes is a mechanism by which cells and enveloped viruses facilitate entry into a new domain. Cell-cell fusion results in the mixing of separate cellular contents into a syncytial cytoplasm, and virus-cell fusion leads to entry of viral genomes into cells. Membrane fusion is directly mediated by fusogens, which are a class of proteins that function to remodel membranes through diverse structural and biochemical mechanisms (Brukman et al., 2019; Segev et al., 2018). Given their function to drive entry of a virus into a cell, viral fusogens are employed in heterologous systems to control delivery of genetic or cellular material to cells. Indeed, VSV-G is the fusogen on vesicular stomatitis virus (VSV) and is utilized on other enveloped viruses to transduce a broad array of cell types in laboratory and clinical settings (Cronin et al., 2005; Milone and O’Doherty, 2018). This process is called pseudotyping and has been used with structurally conserved fusogens from viruses, plants, and worms, to study fusion mechanisms and optimize viral targeting and transduction efficiencies (Avinoam et al., 2011; Nie et al., 2020; Valansi et al., 2017). Whether mammalian cell-cell fusogens that exhibit minimal homology to fusogens in other systems could substitute for viral fusogens and mediate fusion between viruses and cells is unknown.

Myomaker and Myomerger (also known as Myomixer/Minion) are the fusogens for skeletal muscle development and regeneration. Expression of these proteins is spatially and temporally controlled in muscle, where they are not present in quiescent satellite cells but activated during differentiation and then down-regulated in mature myofibers. Myomaker and Myomerger are integral membrane proteins necessary for myoblast fusion, and their expression induces fusion in normally non-fusing cells (Bi et al., 2017; Millay et al., 2013; Quinn et al., 2017; Zhang et al., 2017). However, Myomaker and Myomerger do not possess long extracellular domains typical of classical fusogens that could interact with the trans membrane. Instead, Myomaker and Myomerger function on the cis membrane to modify the local environment and drive fusion through a unique mechanism (Leikina et al., 2018; Millay, 2022). In addition, there are numerous cellular processes, including cell contact and actin nucleation, that cooperate with the myogenic fusogens to drive membrane coalescence in muscle cells (Kim and Chen, 2019). Due to these multiple distinguishing characteristics, it is not intuitively obvious if the myogenic fusogens can function in systems devoid of parallel cellular activity. One barometer to evaluate the fusogenicity by Myomaker and Myomerger is to employ a cell-free system, such as engineered membranes, and assess their ability to replace viral fusogens.

Since the muscle fusogens function specifically in skeletal muscle, their inclusion on viral membranes may target them to muscle. If Myomaker and Myomerger could modify tropism of enveloped viruses to skeletal muscle, it could aid in the delivery of therapeutic modalities. Correction of genetic muscle diseases, such as the muscular dystrophies, is a unique problem due to the distribution of muscle throughout the body in inaccessible locations (Duan et al., 2021). Current approaches include gene restoration by Adeno-associated virus (AAV) (Crudele and Chamberlain, 2019; Tabebordbar et al., 2021), which exhibits tremendous efficacy in pre-clinical models but has not yet achieved full clinical utility. The development of therapeutic enveloped viruses has not progressed even though they possess desirable features including a larger packaging capacity, integration into genomes, transduction of stem cells, and potential for re-dosing (Bulcha et al., 2021). Lentiviral vectors pseudotyped with viral fusogens can transduce dystrophic myofibers and muscle satellite cells, but the efficiency is relatively low and systemic delivery has been difficult due to an immune response against the viral pseudotypes (Gregory et al., 2004; Kimura et al., 2010; Kobinger et al., 2003; Li et al., 2005; MacKenzie et al., 2005). The inability to target enveloped viruses specifically and efficiently to skeletal muscle remains the main obstacle for this class of delivery vehicle.

Here, we establish that Myomaker and Myomerger can drive fusion of basic membranes without the full complement of cellular machinery. As a result, we report the generation of a specialized class of enveloped viruses where Myomaker and Myomerger can functionally substitute for native viral fusogens and direct infectivity specifically towards myogenic targets for delivery of therapeutic material.

## Results

### Myomaker and Myomerger can be functionally pseudotyped on enveloped viruses and alter tropism

To investigate if Myomaker and Myomerger could drive fusion of viral membranes, we utilized a mutant form the vesicular stomatitis virus (VSV), a commonly used pseudotyping platform to assess the fusogenic function of a given protein (Avinoam et al., 2011; Whitt, 2010). The VSV genome was engineered with GFP (serving as a reporter for viral transduction) instead of the gene for the native G protein, which is responsible for viral membrane fusion (VSVΔG) (Figure S1A). Any fusion capability of VSVΔG must therefore be provided in trans by the membrane from the viral producing cells (Figure S1A) (Avinoam et al., 2011; Whitt, 2010). Studies using this system have shown that non-viral fusogens such as the *Caenorhabditis elegans* epithelial fusion failure 1 (EFF-1) and anchor cell fusion failure (AFF-1) proteins, which exhibits structural similarities to viral fusogens, can be used to generate infectious virions (Avinoam et al., 2011; Valansi et al., 2017). To facilitate incorporation of Myomaker and Myomerger on VSVΔG, we transduced BHK21 cell lines with either a control empty vector or vectors for both muscle fusogens, which can be regulated in a doxycycline-dependent manner (Figure S1B). We verified incorporation of the fusogens on viral membranes by immunoblotting (Figure S1C). VSVΔG virions from empty viral producing cells (Bald-VSVΔG) and Myomaker (Mymk) + Myomerger (Mymg) pseudotyped VSVΔG virions (Mymk+Mymg-VSVΔG) were first incubated with VSV-G antibodies to neutralize any remaining VSV-G activity then viruses were applied to BHK21 target cells that harbored an empty vector or expression cassettes for Myomaker and Myomerger. No evidence of transduction was evident by Bald-VSVΔG on empty BHK21 cells although a non-statistically significant increase in GFP^+^ cells was observed in recipient cells expressing Myomaker and Myomerger (Figure S1D and S1E). Mymk+Mymg-VSVΔG virions transduced BHK21 cells expressing Myomaker and Myomerger but not cells with empty vector (Figures S1D and S1E). Mymk+Mymg-VSVΔG also transduced primary cultured mouse myotubes but not proliferating myoblasts or fibroblasts (Figure S1F). These data indicate that Mymk+Mymg-pseudotyped VSVΔG exhibits specific tropism towards cells expressing the muscle fusogens (myotubes and Myomaker + Myomerger expressing BHK21 cells).

Although we have demonstrated successful functional pseudotyping with Myomaker and Myomerger using the VSVΔG system, our ultimate goal is to exploit this hybrid technology towards engineering a safe and efficient muscle-targeting therapeutic vehicle. With multiple pitfalls precluding the use of VSV in a therapeutic setting most notably target cell toxicity (Kopecky et al., 2001) we utilized genome-integrating lentiviruses (LVs) (Figure 1A), another type of enveloped viruses but with greater clinical potential. We generated viral producing HEK293T cells with inducible expression of Myomaker and Myomerger (Figure 1B), and validated the presence of the muscle fusogens, and other viral components, on lentivirus particles produced from these cells (Figures 1C and 1D). Transmission electron microscopy (TEM) analysis revealed relatively uniform rounded morphology of VSV-G pseudotyped lentivirus particles, and similar morphology in Mymk+Mymg virions although some extensions of the viral membranes were observed (Figure 1E) We next tested the transduction capacity of LVs pseudotyped with the muscle fusogens on empty and Myomaker + Myomerger-expressing BHK21 cells. Mymk+Mymg-LVs specifically transduced BHK21 cells expressing Myomaker and Myomerger, in contrast to the broad, non-specific tropism of VSV-G-LVs (Figure 1F). Titers obtained for Mymk+Mymg-LVs were comparable to VSV-G-LV titers (Figure 1G). We observed transduction by Mymk+Mymg-LVs on myotubes (∼80%), low but statistically significant transduction in proliferating myoblasts, but no transduction in fibroblasts (Figure 1H). These findings demonstrate the feasibility for Myomaker and Myomerger pseudotyping on enveloped viruses, including those with therapeutic potential.

**Figure 1.**
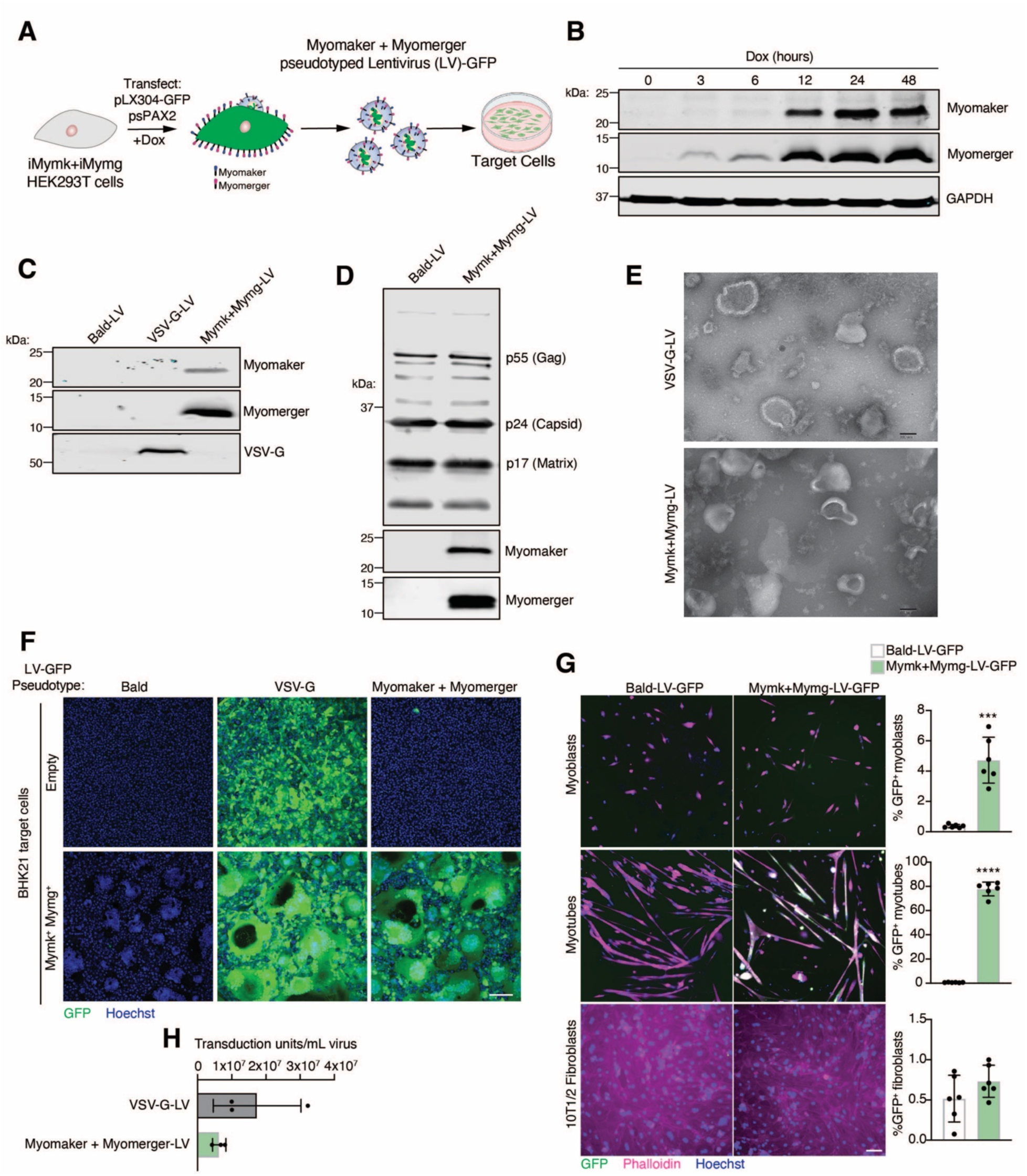
Engineering and characterization of lentiviruses pseudotyped with Myomaker and Myomerger. (A) Schematic showing production of lentiviruses (LV) from Myomaker- and Myomerger-expressing HEK293T cells. (B) Western blot for Myomaker, Myomerger, and GAPDH on HEK293T cell lysates after various times of doxycycline treatment. (C) Western blot on concentrated viral particles for Myomaker, Myomaker, and VSV-G. (D) Western blot on Bald-LVs and Mymk+Mymg-LVs for viral components and the muscle fusogens. (E) Representative images from transmission electron microscopy on VSV-G-LVs and Mymk+Mymg-LVs. Scale bar, 100 nm. (F) Representative images of empty and Mymk+Mymg target cells after application of Bald GFP-encoding LVs or LVs pseudotyped with VSV-G or Myomaker and Myomerger. Nuclei were stained with Hoechst. Scale bar, 100 μm. (G) Quantification of titers for VSV-G-LVs and Mymk+Mymk-LVs prior to concentration of supernatants. Each sample is an average of 3-4 replicates from independent viral preparations. Titers were determined on Mymk+Mymg BHK21 cells. (H) Representative images from cultures of differentiated myotubes, proliferating myoblasts, and fibroblasts that were treated with Bald-LV-GFP or Mymk+Mymg-LV-GFP. Cells were stained with Phalloidin and Hoechst. Scale bar, 100 μm. GFP^+^ cells were quantified for each cell type and histograms are shown on the right. Data are presented as mean ± standard deviation. Statistical test used was an unpaired t-test with Welch’s correction; ***p<0.001, ****p<0.0001.

### Myomaker and Myomerger are independently functional on viral membranes

In our initial efforts, we utilized both muscle fusogens on the viruses and target cells since bilateral expression yields maximum cell-cell fusion efficiency. However, it was not clear if our muscle fusogen virus-cell system is similar to cell-cell fusion where Myomaker is required on both cells and Myomerger is needed on one cell (Quinn et al., 2017; Zhang et al., 2020). Elucidation of the minimal fusogen machinery on the virus and target cells could yield an optimal delivery vehicle and enable cell-type specific tropism. To assess requirements on the virus, we generated viruses (LVs and VSVΔGs) and target cells that contained no fusogen (Empty), Myomaker alone, Myomerger alone, or Myomaker and Myomerger. Virions were produced at comparable levels for each pseudotype based on immunodetection of viral proteins from viral lysates and both Myomaker and Myomerger were independently incorporated on the virus without the need for the other fusogen (Figure 2A). LVs and VSVΔGs with one of the fusogens transduced Mymk+Mymg-expressing target cells at similar efficiencies, but there is an additive effect when both proteins were present on the viral membrane (Figures 2B, 2C, and S2). The ability for enveloped viruses pseudotyped with only one of the fusogens to induce virus-cell fusion suggests that both proteins are making independent contributions on the virus. This is consistent with our previous finding where Myomaker and Myomerger function independently to make distinct contributions to the myoblast fusion reaction (Leikina et al., 2018).

**Figure 2.**
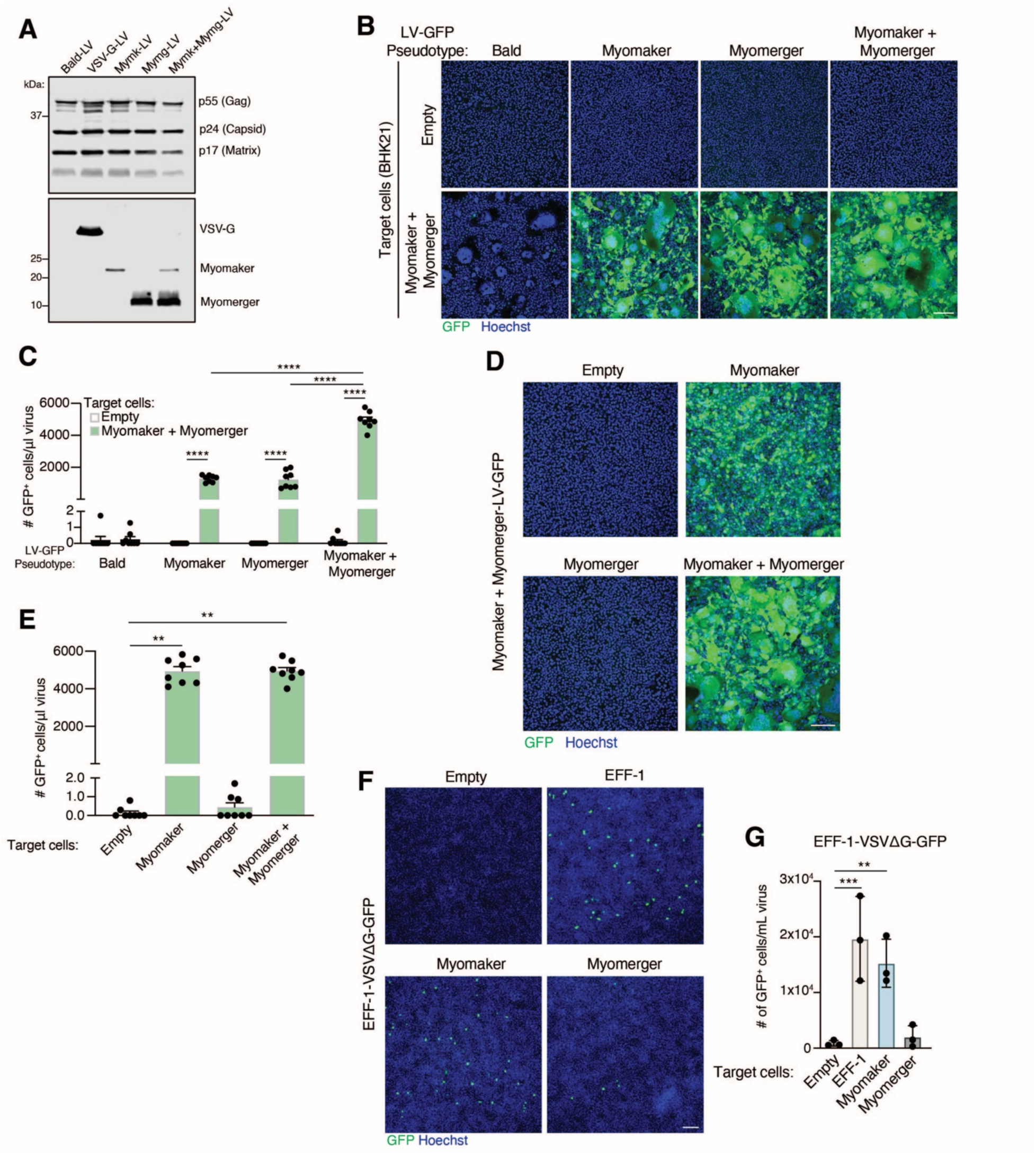
The contribution of Myomaker and Myomerger on viral and cell membranes for pseudotyping. (A) Western blot analysis of viral lysates for the indicated proteins. (B) Representative images for GFP^+^ cells, as an indication of fusion between the indicated lentiviral (LV) pseudotypes and either empty or Myomaker and Myomerger BHK21 target cells. Scale bar, 100 μm. (C) Quantification of viral infection from (B) expressed as # of GFP^+^ cells/μl of virus and assayed from viral supernatants. Pseudotypes with Myomaker alone or Myomerger alone exhibit similar transduction, while an increase is observed with pseudotypes with Myomaker and Myomerger together. (D) Lentiviruses pseudotyped with Myomaker and Myomerger were placed on BHK21 target cells expressing one or both fusogens. Representative images for GFP^+^ cells are shown. Scale bar, 100 μm. (E) Quantification of GFP^+^ cells from (D) as a metric for viral transduction. (F) VSVΔG-GFP pseudotyped with EFF-1 was applied to the indicated BHK21 target cells. EFF-1-VSVΔG-GFP exhibited similar transduction of target cells expressing EFF-1 or Myomaker. Scale bar, 100 μm. (G) Quantification of viral infection from (F) from viral supernatants (prior to concentration). Data are presented as mean ± standard deviation. Statistical tests used were (C) Two-way ANOVA with Tukey’s correction for multiple comparisons; (E) One-way ANOVA (Kruskal-Wallis test) with Dunn’s correction for multiple comparisons; (G) Repeated measures one-way ANOVA with Tukey’s correction for multiple comparisons; **p<0.01, ***p<0.001, ****p<0.0001.

### Myomaker in target cells is sufficient for fusion with viruses containing diverse pseudotypes

To evaluate the requirements on target cells, Mymk+Mymg-pseudotyped LVs or VSVΔGs were applied to empty, Myomaker, Myomerger, or Myomaker and Myomerger BHK21 cells. Target cells expressing Myomaker or Myomaker and Myomerger, but not Myomerger alone, were transduced by Mymk+Mymg-pseudotyped viruses indicating a necessity for Myomaker on target cells (Figures 2D, 2E and S2). To test if Myomaker on target cells increases the general receptivity to viruses, we utilized VSVΔG pseudotyped with EFF-1, which requires an interaction with EFF-1 or a structurally similar fusogen, such as HAP2 from plants, on target cell membranes for infection (Avinoam et al., 2011; Valansi et al., 2017). We confirmed that EFF-1-VSVΔG transduced EFF-1 and not empty target cells, and we also observed transduction by EFF-1-VSVΔG of Myomaker target cells (Figures 2F and 2G). Thus, Myomaker on recipient cells affords as high of a transduction efficiency as EFF-1, the cognate partner. However, virions pseudotyped with the muscle fusogens failed to transduce EFF-1-BHK21 target cells (Figure S2), suggesting that Myomaker activity on the target cells to increase transduction of EFF-1-VSVΔG is not through a physical interaction between Myomaker and EFF-1 as that should also occur when Myomaker is on viruses and EFF-1 on target cells. These data are consistent with a critical function for Myomaker on the target cell for viral receptivity in this specialized system.

### Myomaker and Myomerger pseudotyped lentivirus transduces activated skeletal muscle

We next investigated if LVs pseudotyped with the muscle fusogens are capable of transducing myogenic cells in vivo. Since virus pseudotyped with both muscle fusogens demonstrated maximum transduction efficiency in vitro, we generated Mymk+Mymg-pseudotyped lentivirus encoding Cre (Mymk+Mymg-LV-Cre). Viral supernatants were concentrated and 10^8^-10^9^ virions were injected into the tibialis anterior (TA) muscle of *Rosa*^tdTomato^ mice, which contains a Cre-dependent tdTomato cassette and serves as a readout for viral transduction (Figure 3A). The TAs of these mice were either uninjured or injured by cardiotoxin (CTX) four days prior to injection of the virus (Figure 3A). Two weeks following injection of Mymk+Mymg-LV-Cre, tdTomato^+^ myofibers were observed in CTX-injured, but not in uninjured, muscle (Figure 3B). We also tested transduction in the synergistic ablation model (Goh and Millay, 2017), which causes an increase in muscle load leading to induction of hypertrophy, and is comparatively less damaging than CTX injury but still induces fusion of muscle satellite cells (Figure 3C). Here, we also utilized a Bald-pseudotyped lentivirus encoding Cre as a control and similar levels of virions were produced compared to Mymk+Mymg-LVs (Figure 3D). TdTomato^+^ myofibers were only detected in the plantaris muscle after muscle overload and injection with Mymk+Mymg-LV-Cre (Figure 3D). These data indicate that lentiviruses pseudotyped with Myomaker and Myomerger can transduce wild-type muscle in vivo when the muscle fusogens are expressed, such as following injury or a hypertrophic stimulus.

**Figure 3.**
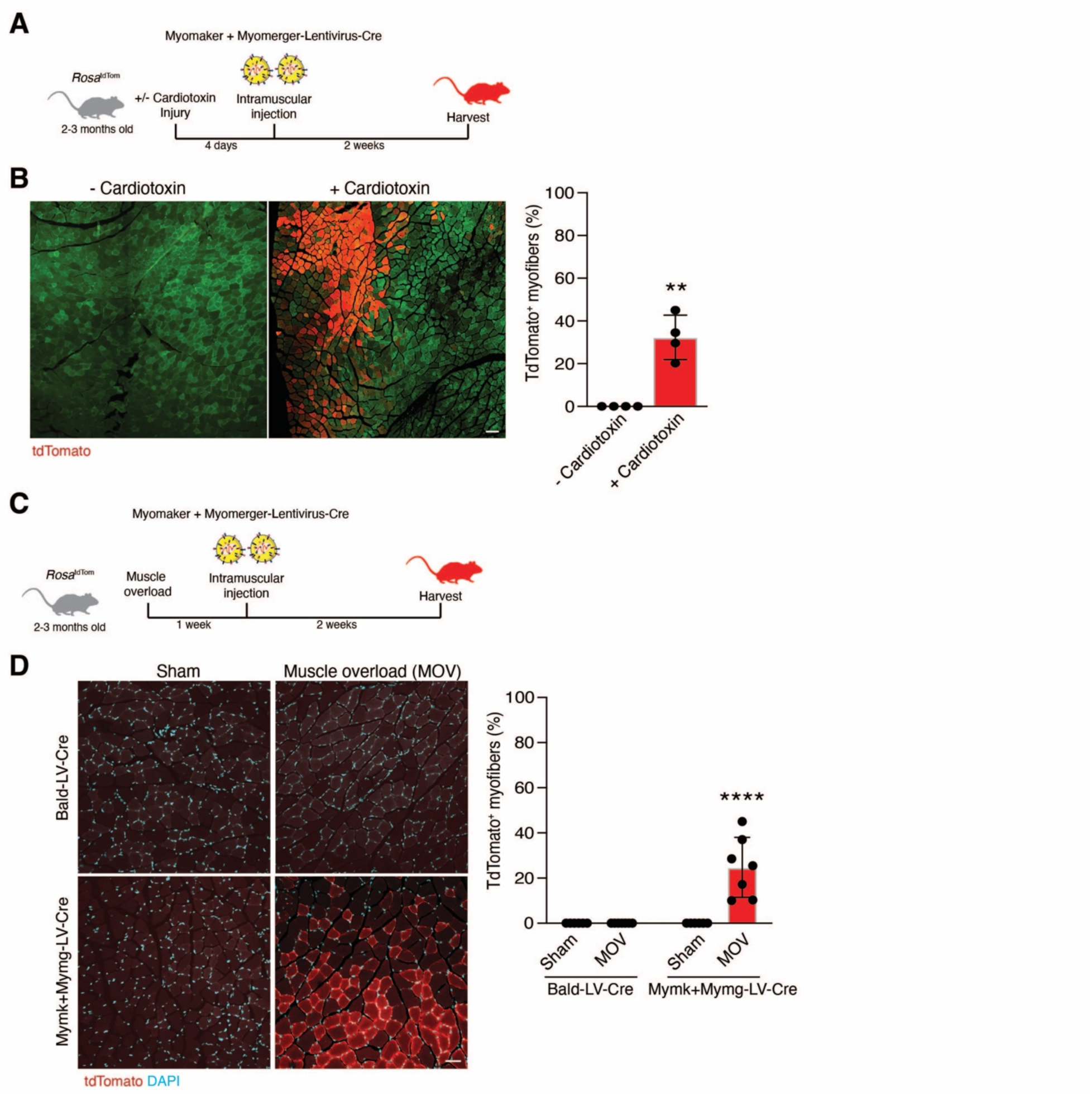
Mymk+Mymg-LVs transduce activated skeletal muscle in vivo. (A) Lentiviruses encoding for Cre recombinase and pseudotyped with Myomaker and Myomerger (Mymk+Mymg-LV-Cre) were produced and delivered to the tibialis anterior muscles of *Rosa*^tdTom^ mice through direct intramuscular injection. Some tibialis anterior muscles were injured with cardiotoxin prior to receiving lentivirus. (B) Representative images are shown displaying tdTomato^+^ myofibers after injury and regeneration. Green is muscle autofluorescence. The percentage of tdTomato^+^ myofibers is shown (right panel). Scale bar, 100 μm. (C) Similar setup as in (A) except muscle overload, which causes hypertrophy, was performed and lentivirus was injected into the plantaris. (D) Representative images are shown for sham and overloaded muscles treated intramuscularly with Bald-LV-Cre or Mymk+Mymg-LV-Cre. Scale bar, 50 μm. Quantification of the percentage of tdTomato^+^ myofibers (right panel). Data are presented as mean ± standard deviation and combined from at least two independent lentiviral preparations. Statistical tests used were (B) unpaired t-test with Welch’s correction; (D) Two-way ANOVA with Tukey’s correction for multiple comparisons; **p< 0.01, ****p< 0.0001.

We next applied the Mymk+Mymg-LV technology to *mdx*^4cv^ mice, a model of Duchenne muscular dystrophy, which is a devastating genetic muscle disease resulting in chronic cycles of muscle injury and regeneration (O’Brien and Kunkel, 2001). Bald- or Mymk+Mymg-LVs encoding Cre (10^8^-10^9^ virions) were injected into the TA muscles of *mdx*^4cv^; *Rosa*^tdTomato^ mice and viral transduction was assessed two weeks later (Figure 4A). Large areas of tdTomato^+^ myofibers were detected in muscles injected with Mymk+Mymg-LV-Cre without prior CTX injury, and the percentage of transduced myofibers increased after CTX (Figure 4B). Since we observed an increase in tdTomato^+^ myofibers in settings when myogenic progenitors are activated (injury, muscle overload, dystrophic muscle), we postulated that Mymk+Mymg-LVs may be trophic for myogenic progenitors and assessed tdTomato expression in α7-Integrin^+^ myogenic cells five weeks following intramuscular injection of pseudotyped LVs in TA muscles of *mdx*^4cv^; *Rosa*^tdTomato^ mice. Standard FACS analysis was used to identify the α7-Integrin^+^ cells (Figure S3) (Maesner et al., 2016), and revealed that ∼50% of α7-Integrin^+^ cells were positive for tdTomato following Mymk+Mymg-LV-Cre but not Bald-LV-Cre administration (Figure 4C). Indicating specificity of Mymk+Mymg-LVs for myogenic cells, the number of tdTomato^+^ non-myogenic cells was negligible (Fig. 4C). In a separate experiment, we generated Mymk+Mymg-pseudotyped lentivirus encoding luciferase (Luc) and compared its transduction capacity to VSV-G-pseudotyped lentivirus following intramuscular injection of CTX-injured *mdx*^4cv^ TA muscles (Figure 4D). Luciferase activity was significantly higher in TA muscles injected with Mymk+Mymg-LV-Luc as compared to that injected with VSV-G-LV-Luc one week following viral injections (Figure 4D). Moreover, time course analysis revealed an increase in reporter signal in Mymk+Mymg-LV-Luc treated animals over time (Figure 4D). We interpret the increase in luciferase signal over time to be contributed by the ongoing fusion of transduced muscle satellite cells. Taken together, these results highlight that after local injection, Mymk+Mymg-pseudotyped virus can transduce dystrophic skeletal muscle cells including myogenic progenitors.

**Figure 4.**
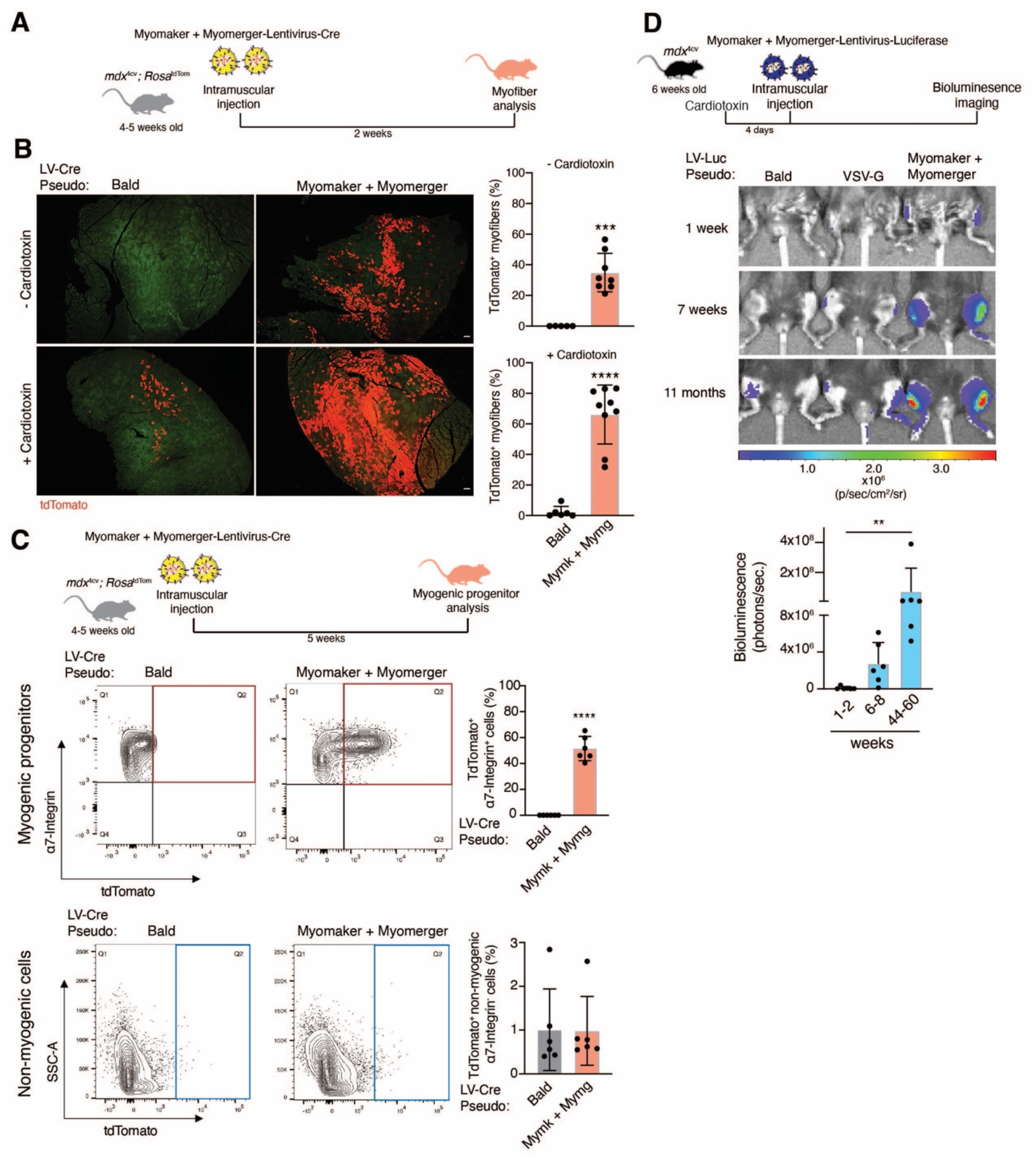
Myofibers and myogenic progenitors in dystrophic mice are transduced by lentiviruses pseudotyped with Myomaker and Myomerger. (A) *Mdx*^4cv^; *Rosa*^tdTom^ mice were used as recipients for Bald-LV-Cre or Mymk+Mymg-LV-Cre. Tibialis anterior muscles receiving lentiviruses through intramuscular injection were uninjured or previously injured with cardiotoxin. (B) Representative images showing tdTomato^+^ myofibers with and without cardiotoxin injury. Green is muscle autofluorescence. Scale bar, 100 μm. Quantification of the percentage of tdTomato^+^ myofibers (right panel). (C) Top panel displays experimental design for analysis of muscle progenitors. Middle panels show representative FACS plots and quantification for α7-Integrin^+^ myogenic cells (y-axis) and tdTomato^+^ cells (x-axis) from muscle that received Bald-LV-Cre or Mymk+Mymg-LV-Cre. Bottom panels show representative FACS plots and quantification for tdTomato^+^ non-myogenic interstitial cells. (D) Top panel is the experimental design using lentivirus encoding for luciferase that was pseudotyped with Bald, VSV-G, or Myomaker and Myomerger. After cardiotoxin injury, lentiviruses were injected intramuscularly and bioluminesence measured through IVIS imaging multiple time points after viral delivery. Representative pseudocolored images show a progressive increase in bioluminescence in the muscles treated with Myomaker and Myomerger-pseudotyped virus. Bottom panel is quantification of bioluminescence for muscles transduced with lentivirus containing Myomaker and Myomerger. Data are presented as mean ± standard deviation and combined from at least two independent lentiviral preparations. Statistical tests used were (B) Unpaired t-test with Welch’s correction; (C) unpaired t-test with Welch’s correction; (D) Friedman test with Dunn’s correction for multiple comparisons; **p<0.01, ***p<0.001, ***p< 0.0001.

### Myomaker and Myomerger pseudotyped lentivirus specifically transduces skeletal muscle following systemic delivery

An important consideration for a delivery vehicle is an ability to specifically transduce muscle after systemic delivery. We assessed transduction of non-muscle and skeletal muscle tissues through evaluation of tdTomato labeling following systemic delivery of Bald- or Mymk+Mymg-LV-Cre (10^8^-10^9^ virions) to *mdx*^4cv^; *Rosa*^tdTomato^ mice. LVs were administered to mice through three separate retro-orbital (RO) injections starting at two-week of age and multiple tissues were analyzed two weeks after the final injection (Figure 5A). Previous reports have indicated minimal VSV-G-LV transduction in most tissues including skeletal muscle following systemic delivery likely due to its neutralization by an activated immune response towards VSV-G (Li et al., 2005). In contrast, we observed viral transduction in various skeletal muscles by Mymk+Mymg-LV-Cre but not Bald-LV-Cre (Figures 5B and 5C). Notably, we observed maximum and robust transduction in the diaphragm muscle (∼80% of myofibers) (Figures 5B and 5C), which is critical to target since respiratory failure is a major reason for mortality in many skeletal myopathies (Wahlgren et al., 2022). We observed transduction of approximately 10% of myofibers in the limb muscles (Figure 5C). Moreover, we did not detect evidence of viral transduction in non-skeletal muscle tissues including heart, kidney, liver, and spleen (Figure 5D), highlighting the specificity of this pseudotyped vehicle. The lack of transduction in non-muscle tissues is expected given the absence of Myomaker and Myomerger expression and cellular fusion in those tissues. We also treated *Rosa*^tdTomato^ mice with Bald-LV-Cre and Mymk+Mymg-LV-Cre through a single retro-oribital injection during muscle overload, and detected tdTomato^+^ myofibers in muscle undergoing hypertrophy from mice that were treated with Mymk+Mymg-LV-Cre (Figures 5E and 5F). Altogether, these findings are supportive of a skeletal muscle-specific targeting role for systemically delivered Mymk+Mymg-pseudotyped viruses in the dystrophic myopathies and during muscle overload.

**Figure 5.**
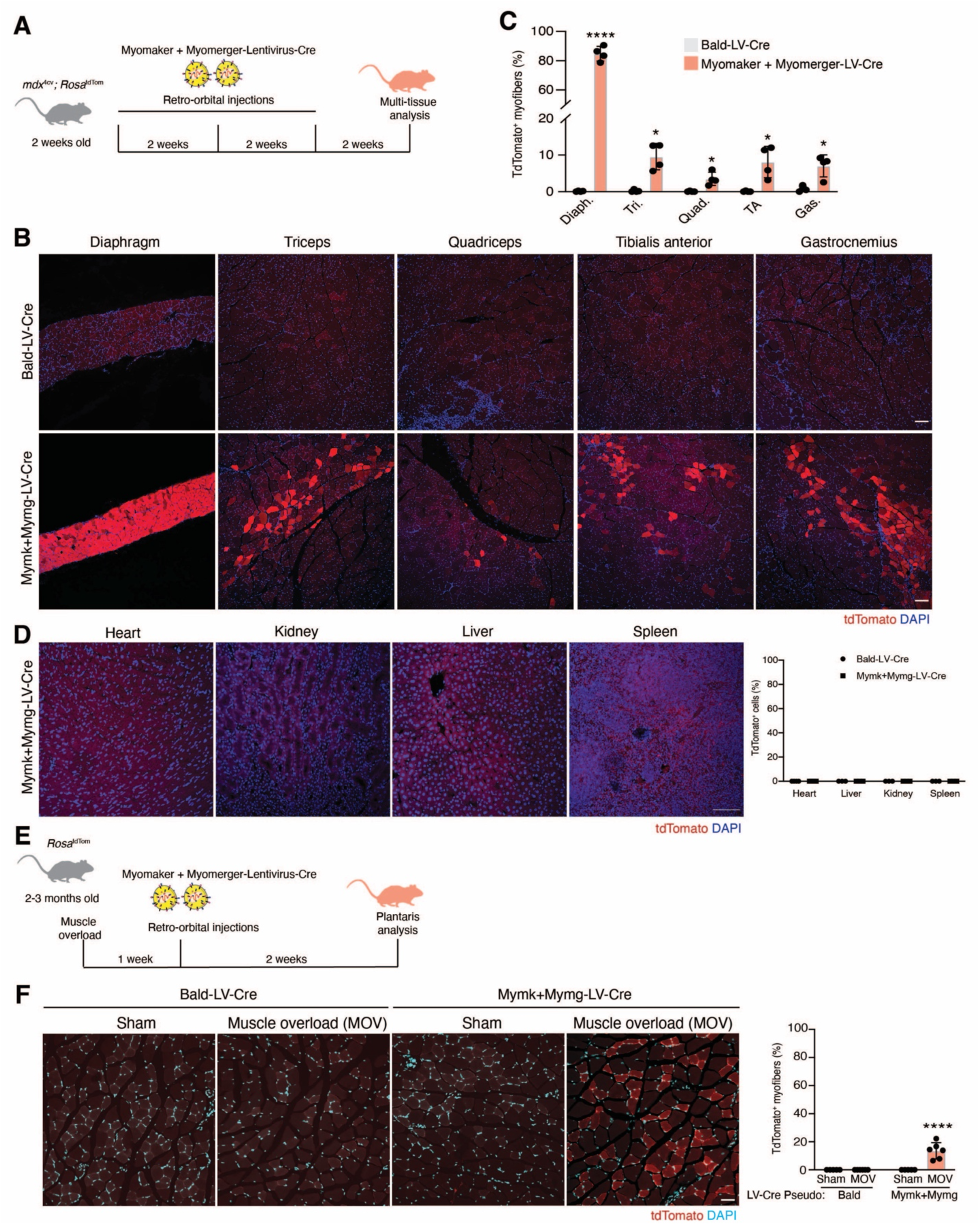
Tropism of Myomaker+Myomerger lentiviruses is specific for skeletal muscle after systemic delivery. (A) Schematic of experimental plan to assess transduction of lentiviruses delivered to *mdx*^4cv^; *Rosa*^tdTom^ mice through three retro-orbital injections two weeks apart. (B) Representative images showing tdTomato^+^ myofibers from various skeletal muscle tissues after retro-orbital delivery of Cre-encoding lentiviruses pseudotyped with Bald or Myomaker and Myomerger. Scale bar, 100 μm. (C) Quantification of the percentage of tdTomato^+^ myofibers. (D) Representative images analyzing tdTomato^+^ cells from non-skeletal muscle tissues. No positive cells were detected. Quantification is shown on the right. Scale bar, 100 μm. (E) Experimental plan for systemic delivery of lentiviruses after muscle overload. (F) Representative images showing tdTomato^+^ myofibers in the plantaris. The percentage of tdTomato^+^ myofibers is also shown (right panel). Scale bar, 50 μm. Data are presented as mean ± standard deviation and combined from at least two independent lentiviral preparations. Statistical tests used were (C) Unpaired t-tests with Welch’s correction; (F) Two-way ANOVA with Tukey’s correction for multiple comparisons; *p<0.05, ****p<0.0001.

### Myomaker and Myomerger pseudotyped lentivirus delivers therapeutic material to dystrophic muscle and alleviates myopathy

We next assessed the ability of Mymk+Mymg-LVs to deliver therapeutically relevant material. We packaged a miniaturized form of the dystrophin gene (μDys5) that can partially compensate for full-length dystrophin (Ramos et al., 2019), in Bald-LVs or Mymk+Mymg-LVs (Figure 6A). Viral particles were injected into TA muscles of *mdx*^4cv^ mice and analyzed 5 weeks later for μDys by immunohistochemistry (Figure 6B). Consistent with the previous functional readout of reporter genes packaged by Mymk+Mymg-LVs in this study, we observed a wide distribution of myofibers expressing μDys while no such expression was detected in muscle treated with Bald-LVs (Figures 6B and 6C). We also tested if μDys could be delivered by Mymk+Mymg-LVs through three doses of systemic retro-orbital injections (Figure 6D). For mice treated with Mymk+Mymg-LV-μDys, μDys was detected in 5-25% of myofibers in limb muscles (Figure S4) and 77-90% of myofibers in the diaphragm (Figure 6E). Mymk+Mymg-LVs delivered enough μDys to the diaphragm to reduce indices of pathology including centrally nucleated myofibers and fibrosis (Figures 6F and 6G). Overall, these results provide proof-of-concept experimental evidence for the therapeutic utility of enveloped viruses pseudotyped with the muscle fusogens.

**Figure 6.**
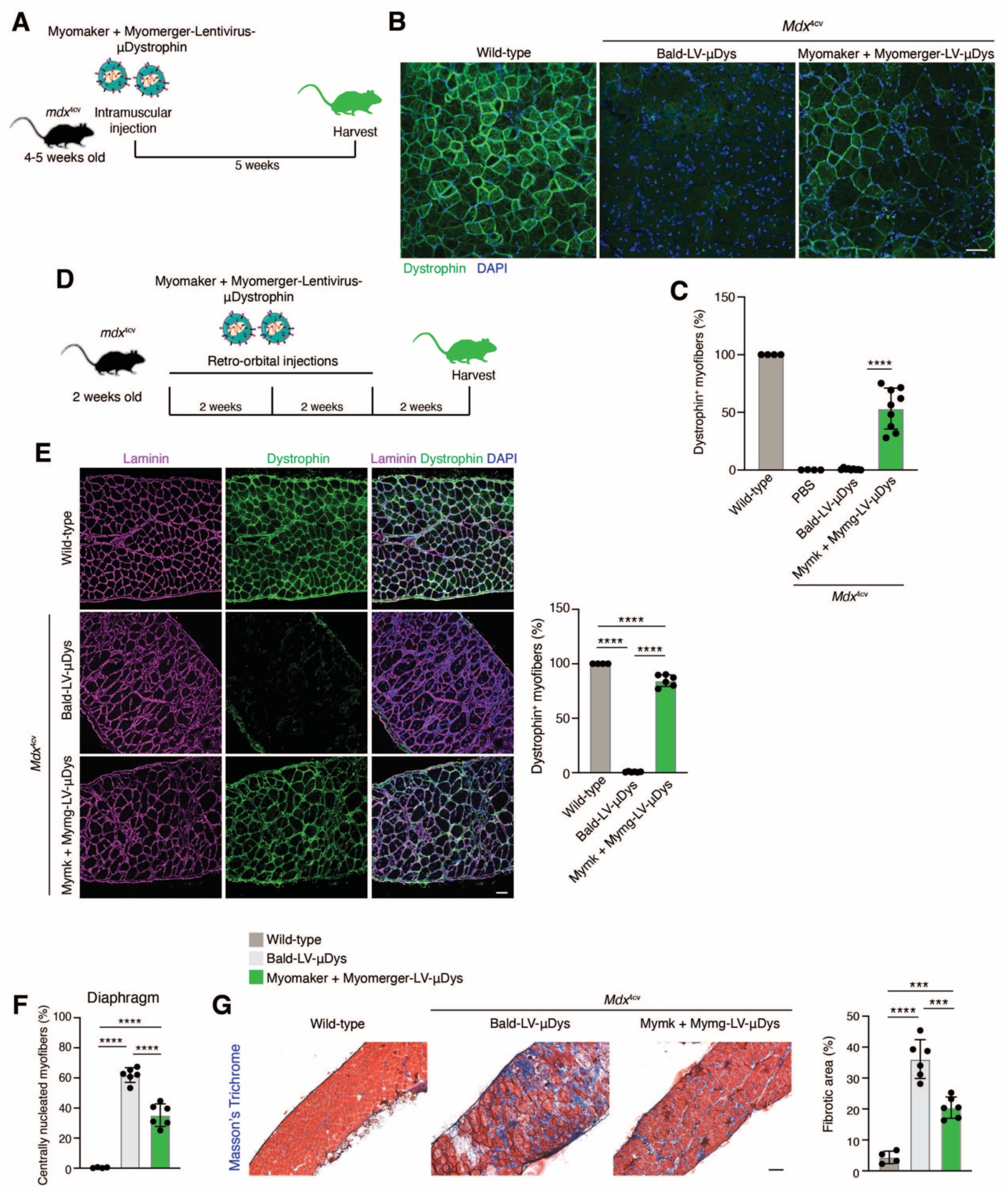
Lentiviruses pseudotyped with the muscle fusogens deliver therapeutic material to skeletal muscle. (A) Schematic for experimental plan to assess transduction of Myomaker and Myomerger-pseudotyped lentiviruses packaged with μDystrophin (μDys) after intramuscular injection. (B) Immunostaining for dystrophin from tibialis anterior muscles from wild-type or *mdx*^4cv^ mice that were injected with Bald- or Myomaker and Myomerger-lentiviruses encoding μDys. Scale bar, 100 μm. (C) Quantification of the percentage of μDys^+^ myofibers from (B). (D) Design to assess systemic delivery of lentiviruses in *mdx*^4cv^ mice. (E) Immunostaining for dystrophin in *mdx*^4cv^ diaphragms treated with Bald-LV-μDys or Mymk+Mymg-LV-μDys. Scale bar, 50 μm. Quantification of μDys^+^ myofibers is shown on the right. (F) Quantification of centrally nucleated myofibers in the diaphragms from wild-type mice and mice treated with Bald-LV-μDys or Mymk+Mymg-LV-μDys. (G) Masson’s trichrome staining of diaphragms treated systemically with the indicated lentivirus. Scale bar, 100 μm. Quantification of blue trichrome area is shown on the right. Data are presented as mean ± standard deviation and combined from at least two lentiviral preparations. Statistical tests used were (C), (E-F) One-way ANOVA with Tukey’s correction for multiple comparisons; ***p<0.001, ****p<0.0001.

### Myomaker and Myomerger enhance tropism of extracellular vesicles to muscle cells

Since Myomaker and Myomerger endowed tropism of viral membranes for skeletal muscle cells, we asked if this effect was generalizable to other types of membrane-derived therapeutic vectors. We collected extracellular vehicles (EVs) (van Niel et al., 2018) from 10T½ fibroblasts that were engineered to overexpress Myomaker, Myomerger, or Myomaker and Myomerger (Figure S5A). Western blot analysis showed the presence of Myomaker and Myomerger on EVs (Figure S5B). EVs were labeled with the lipid probe DiI and placed on myotubes, where we observed significantly more uptake of EVs that contained one or both muscle fusogens (Figures S5C and S5D). These data show that the skeletal muscle fusogens could enhance tropism of non-viral membrane vectors to skeletal muscle cells.

## Discussion

The muscle-specific proteins Myomaker and Myomerger drive the final steps of skeletal muscle cell fusion. The work described in this study addressed two main questions: a) can Myomaker and Myomerger drive fusion of cell-free membranes and if so, b) can the muscle-specific fusogens direct tropism of non-cellular membranes towards myogenic cells. These questions needed to be investigated and answered because it was unclear whether the functional properties of the muscle fusogens to modulate the cis membrane could be transferred to non-cellular membranes like viral membranes and substitute for native viral-fusogens that act on the trans membrane. The pseudotyping results shown here advance our understanding of these fusogens as they confirm the ability of Myomaker and Myomerger to function in the absence of additional cellular processes implicated in muscle cell-cell fusion. We believe this pseudotyping platform, where the individual activity of each of the muscle fusogens can be isolated will be utilized in future studies to better understand cell fusion mechanisms. Development of viral envelopes that are trophic for skeletal muscle as shown here has therapeutic relevance because it has the potential to complement the limitations of current viral-based gene therapies.

A major issue with gene and cell therapies is an ability to deliver material specifically to the tissue of interest. We establish that vectors pseudotyped with the muscle fusogens are highly specific for skeletal muscle cells after local and systemic delivery. In vitro, transduction is only observed when Myomaker is present in target cells and in vivo transduction by Mymk+Mymg-LVs is restricted to skeletal muscle conditions associated with activated muscle progenitors that express the muscle fusogens. These data overall suggest that the cell type in muscle that is transduced in vivo is one which expresses the muscle fusogens, and at a minimum Myomaker. In dystrophic muscle, Myomaker is expressed in activated muscle progenitors derived from satellite cells and regenerating myofibers that exhibit Myomaker expression contributed from the fusion of progenitors (Petrany et al., 2020). The need for Myomaker in target cells explains why uninjured wild-type muscle is not transduced since there are no Myomaker-expressing activated progenitors or regenerating myofibers. Targeting activated muscle progenitors raises the possibility that this pseudotyping approach affords the opportunity to permanently modify the resident satellite cell population, which could offer a supply of therapeutically corrected cells that can continue to fuse with myofibers over the lifetime of the recipient. Indeed, the presence of transduced muscle progenitors five weeks after intramuscular viral delivery and the increase in luciferase signal in myofibers after 11 months strongly suggest that satellite cells are targeted. Targeting of satellite cells could result from direct transduction by Mymk+Mymg-LVs or transduction of activated progenitors that undergo self-renewal (Cutler et al., 2021).

Systemically injected Mymk+Mymg-LVs into neonatal and adult mice target all skeletal muscle tissues analyzed at varying efficiencies. 5-25% of myofibers in limb muscles are targeted by Mymk+Mymg-LVs two weeks after the final injection, but the potential for Mymk+Mymg-LVs to target progenitor cells could yield a progressive increase in delivery to myofibers through continued fusion over time. Of note, Mymk+Mymg-LVs exhibit an impressive ability to target the diaphragm (77-90% of myofibers), a muscle critical for respiration and often the cause of death for patients with muscle diseases. Consistent with targeting the majority of myofibers in the diaphragm, we observed a reduction in pathologic indices such as reduced central nucleation and fibrosis in dystrophic mice systemically treated with Mymk+Mymg-LVs. AAV9 and Myo-AAV exhibit more robust transduction after a single dose (Ramos et al., 2019; Tabebordbar et al., 2021) because the vector is highly trophic for mature myofibers that are always present in high levels in dystrophic muscle. Detection of μDys expression only after multiple systemic injections, and the variability in targeting efficiencies for different muscle groups, is likely related to the specificity of Mymk+Mymg-LVs for Myomaker-expressing progenitors or regenerating myofibers, the presence of which fluctuate in dystrophic muscle depending on whether the muscle is undergoing degeneration or regeneration.

To the best of our knowledge, this study is the first to report the generation of a viral vector pseudotyped with mammalian fusogens that can function as a therapeutic delivery vehicle for a specific cell-type. Mymk+Mymg-LVs overcome a major obstacle in lentiviral gene therapy for muscle and adds to potential approaches to correct muscle diseases. Lentiviral vectors have been the subject of decades of research (Connolly, 2002; Escors and Breckpot, 2010; Kimura et al., 2010; Milone and O’Doherty, 2018) due to desirable features like genomic integration, relatively large packaging capacity, and transduction of stem cells which makes them a promising vector for potential lifelong supply of an absent or defective protein. Despite these desirable qualities, their therapeutic application has been hindered due to a reliance on viral pseudotypes, such as VSV-G, that are neutralized by the complement system upon systemic delivery (DePolo et al., 2000). Another general concern for LV-based therapy is oncogenesis due to random insertion into the genome (Schlimgen et al., 2016; Themis et al., 2005), but this may not be an issue since myofibers are a multinucleated terminally differentiated cell type and there are methods to bias integration (Cai et al., 2016). Thus, the unique advantages of Mymk+Mymg-LVs have the potential to complement some enduring challenges associated with AAV gene therapy, which currently dominates gene-corrective strategies for muscle. AAV gene therapy has potential issues including pre-existing immunity, toxicity, off-target effects, dosing-limitations, and limited transgene persistence (Duan, 2018; Manini et al., 2021; Morales et al., 2020; Morgan and Muntoni, 2021), indicating that alternative vectors need to be developed.

Here we use a Duchenne muscular dystrophy model as proof-of-concept for the efficacy of Mymk+Mymg-LVs to deliver therapeutically relevant genetic material. However, we think that this platform could be utilized for a wide range of genetic and acquired muscle diseases, and even for non-muscle diseases that would benefit from expression of a secreted factor like Factor IX for hemophilia (Buchlis et al., 2012). For the cardiac components of pathologies, such as Duchenne muscular dystrophy, Mymk+Mymg-LVs would need to be used in combination with another therapeutic modality. One could imagine that Mymk+Mymg-LVs could be used to target progenitor cells and regenerating myofibers when the availability of those cell types are high in skeletal muscle, then AAVs could be used to further elevate dystrophin expression in mature myofibers and cardiac cells. For muscle diseases not affecting the heart, such as X-linked myotubular myopathy, Mymk+Mymg-LVs could be a stand-alone therapy especially if patients have antibodies to AAV capsids. Clearly, the precise delivery mechanism for lifelong correction of genetic muscle diseases cannot be currently ascertained. The data presented here shows that Mymk+Mymg-LVs have potential especially considering this is the first description of a delivery vehicle that harnesses the muscle-specific fusion machinery to drive tropism. We expect the foundational therapeutic vehicle described in this study can be further optimized with additional proteins and lipids as mechanistic details of the myoblast fusion reaction are uncovered. Moreover, non-viral membrane vehicles, such as extracellular vesicles containing Myomaker and Myomerger, could be utilized as also shown in this study. Overall, our findings demonstrate that the fusogenic properties of mammalian muscle fusogens can be transferred to viral membranes thus establishing a specialized class of delivery vehicles that are specific for skeletal muscle.

## Acknowledgments

We thank members of the Millay laboratory and N. Scott Blair (Molkentin lab) for reagents and discussion; Victoria Summey of the Cincinnati Children’s Hospital Medical Center Comprehensive Mouse and Cancer Core for help with retro-orbital injections. This work was mainly supported by a grant to D.P.M. from the National Institutes of Health (R61AR076771). Work in the Millay laboratory is also funded by grants to D.P.M. from Children’s Hospital Research Foundation, National Institutes of Health (R01AR068286, R01AG059605), and a sponsored research agreement with Sana Biotechnology. Work in the Podbilewicz laboratory is funded by grants to B.P. from the Israel Science Foundation (grants 257/17, 2462/18, 2327/19, and 178/20). The Chamberlain laboratory is supported by the Sen. Paul D. Wellstone Muscular Dystrophy Specialized Research Center (P50 AR065139).

## Author contributions

S.M.H., B.P., and D.P.M. conceived the project. S.M.H., M.J.P., E.G., L.F., A.A.W.C., and V.P. conducted experiments and analyzed the data. M.A.W., J.S.C., B.P., and D.P.M. supervised the project. S.M.H. and D.P.M. wrote the manuscript with input from all authors.

## RESOURCE AVAILABILITY

### Lead Contact

Please direct request for resources and reagents to Lead Contact: Douglas P. Millay

### Materials availability

Plasmids and cell lines generated in this study are available upon request.

## EXPERIMENTAL MODEL AND SUBJECT DETAILS

### Animals

All mice used in this study were maintained on a C57BL/6 background including C57BL/6 (wild-type), *mdx*^4cv^ (stock #002378; The Jackson Laboratory, Bar Harbor, ME, USA), *Rosa26*^tdTomato^ (stock #007905; The Jackson Laboratory). *Mdx*^4cv^; *Rosa26*^tdTomato^ mice were generated by crossing *mdx*^4cv^ mice with *Rosa26*^tdTomato^ mice. All experiments were performed on gender- and age-matched cohorts, where both male and female mice were used, and animals were randomly assigned to control or experimental groups. All animal procedures were approved by Cincinnati Children’s Hospital Medical Center’s Institutional Animal Care and Use Committee and were conducted in accordance with Association for Assessment and Accreditation of Laboratory Animal Care International guidelines.

### Cells

Primary myoblasts were isolated from wild-type mice as described previously (Hindi et al., 2017). Cells were plated on matrigel-coated plates, maintained in growth medium (20% fetal bovine serum (Peak Serum) and 2.5 ng/ml human bFGF (Gibco) in Ham’s F-10 nutrient mixture (Gibco) with penicillin/streptomycin (Gibco), and differentiated in differentiation media (2% horse serum (Gibco) in high-glucose DMEM (Gibco), with penicillin/streptomycin). 10T½ fibroblasts were purchased from American Type Culture Collection and propagated in DMEM (Gibco) containing 10% heat-inactivated fetal bovine serum and supplemented with antibiotics and sodium pyruvate (Gibco). HEK293T and C2C12 cells were purchased from American Type Culture Collection (ATCC), and Baby hamster kidney (BHK21) cells were provided by Dr. Michael Whitt. Cells were propagated in DMEM (Gibco) containing 10% heat-inactivated fetal bovine serum and supplemented with antibiotics along with sodium pyruvate for HEK293T and BHK21 cells and L-glutamine (Gibco) for BHK21 cells. All doxycycline inducible expression cell lines were propagated in DMEM containing 10% Tet-Free fetal bovine serum (Peak Serum). Platinum-E retroviral packaging cells were purchased from Cell Biotech and propagated under constant selection conditions in DMEM (Gibco) containing 10% heat-inactivated fetal bovine serum and supplemented with antibiotics and sodium pyruvate. All cells were grown and maintained in a 37°C incubator with 5% CO_2._

### Method details

#### Traditional (VSV-G-pseudotyped) lentivirus generation

5x10^6^ HEK293T cells were plated on a 100 mm dish and transfected with a cocktail of 10 μg transfer plasmid, 7.5 μg psPAX2 packaging plasmid (Addgene, plasmid #12260), and 2.5 μg pMD2.G envelope plasmid (Addgene, plasmid #12259) with 60 μl FuGENE-6 (Promega Madison, WI, USA) or polyethylenimine (PEI) (linear, MW 250,000) (Polysciences Warrington, PA, USA) or in 1ml DMEM. For experiments using PEI, media was changed after 4h and supernatant containing viral particles was collected 40-48h later. For viral preparation utilizing FuGENE-6, no media change was performed and supernatant containing viral particles was collected 48h later. Viral supernatants were centrifuged at 2500 rpm for 2.5 minutes and passed through a 0.45 μm SFCA filter before use. For lentivirus utilized in generating viral producing BHK21 and HEK293T cells, mouse Myomaker, mouse Myomerger, EFF1-V5, were cloned into the transfer lentivirus plasmid pLVX-TetOne-Puro (Clontech, plasmid #631849) or pLVX-TetOne-Blast (modified from Clontech, plasmid #631849).

### Retrovirus generation

5x10^6^ Platinum-E retroviral packaging cells (Cell biotech, San Diego, CA, USA) were transfected with a cocktail of 10 μg of pBabe-X-Myomaker and/or pBabe-X-Myomerger, or pBabe-X-empty plasmid along with 30 μl FuGENE-6 in DMEM. After 48 hours, supernatants containing retroviral particles were centrifuged at 2500 RPM for 2.5 minutes and passed through a 0.45 μm SFCA filter before use.

### Myomaker, Myomerger, EFF1-V5 pseudotyped viral producing and recipient cell lines

Baby hamster kidney (BHK21) cells were transduced with lentivirus coding for doxycycline-inducible Myomaker and/or Myomerger, EFF1-V5, or an empty cassette to generate cells that would produce pseudotyped VSV. BHK21 cell lines generated here were also used as viral recipient cells in this study. For generation of lentiviral producing cell lines with various pseudotypes, HEK293T cells were transduced with traditional VSV-G pseudotyped lentivirus coding for doxycycline inducible Myomaker and/or Myomerger, or empty plasmid. Transduction of both BHK21 and HEK293T cells to express the fusogen (Myomaker and/or Myomerger, or EFF1-V5) was achieved through spinfection by centrifugation at 600xg for 1 hour at 22°C immediately following viral overlay and incubated overnight. Cells were then selected with puromycin or blaticidin for 72 hours.

The above BHK21 cells were used as recipient cells for assessment of function and titering. For Cre-coding lentiviral pseudotypes, BHK21 recipient cells were generated by initial transduction with traditional VSV-G lentivirus coding for a Cre-reporter (plasmid #62732, Addgene, Watertown, MA, USA), followed by transduction with empty or doxycycline inducible Myomaker + Myomerger -lentivirus (traditional VSV-G pseudotyped) in a manner as described above.

### Extracellular vesicle producing cells

10T½ fibroblasts were transduced with retrovirus generated from platinum E cells which were transfected with pBabeX-Myomaker and/or pBabeX-Myomerger, or pBabeX-empty. Fibroblasts were subjected to spinfection by centrifugation at 600xg for 1 hour at 22°C immediately following viral overlay and incubated overnight.

### Generation of pseudotyped vesicular stomatitis virus

0.5x10^6^ Myomaker and/or Myomerger, EFF-1-V5, or empty BHK21 cells (all dox-inducible) were plated on a 60 mm cell culture plate with 250 ng/ml doxycycline for 36 hours. Cells were then infected with VSVG-complemented VSVΔG recombinant virus (VSVΔG-G) for 2 hours in serum free DMEM. Virus infected cells were washed 3 times with PBS to remove unabsorbed VSVΔG-G virus. Following a 24-hour incubation, the supernatant containing the VSVΔG-pseudovirus was harvested and centrifuged twice at 2500 RPM for 5 minutes to remove cellular debris. Pseudotyped viral supernatants were then treated with anti-VSV-G (1:1000, #8G5F11, Kerafast) to neutralize any residual VSV-G. Viral supernatants were either immediately used for functional analysis or concentrated by ultracentrifugation for detection of fusogenic proteins on viral particles.

### Generation of lentivirus pseudotyped with the muscle fusogens

2.5x10^6^, 5x10^6^, or 10x10^6^ Myomaker and/or Myomerger, or empty inducible HEK293t cell lines generated in this study were plated on 60-, 100-, or 150-mm dishes respectively. After 24 hours, cells were then treated with 1 μg/ml doxycycline for 2 hours after which media was changed to 100 ng/ml doxycycline in 2% horse serum in DMEM. Cells were then transfected with a cocktail of lentivirus transfer plasmid (5, 10, or 20 μg) carrying GFP, Cre, Luciferase, or μDystrophin (GFP and μDystrophin were inserted into a pLX304 backbone (Addgene, Plasmid #25890), Cre replaced eGFP in pLKO.1-puro eGFP (Sigma Aldrich, plasmid #SHC005) and Luciferase plasmid was directly purchased from Addgene (pLX304 Luciferase-V5 (Addgene, plasmid #98580)) along with lentiviral packaging plasmid (3.75, 7.5, or 15μg psPAX2) and FuGENE-6 (Promega) or PEI (Polysciences) at a ratio of 1:3 μg DNA:μl transfection reagent in DMEM. After 16-24 hours, viral supernatants were collected, cellular debris was cleared, and passed through a 0.45 μm SFCA filter and either immediately used for functional analysis, concentrated, or stored at -80°C for later use. Remaining cells were replenished with fresh 2% horse serum in DMEM supplemented with 50 ng/ml doxycycline for another 16-24 hours where a second collection of viral supernatants was performed and processed as described for the first viral supernatant collection.

### Functional assessment of pseudotyped virus in myogenic and non-myogenic cells

Primary myoblasts, myotubes, or 10T½ fibroblasts were plated on 48-well cell culture plates and overlayed with pseudotyped VSVs or LVs coding for GFP with 6 μg/ml polybrene and subjected to spinfection as described above. Viruses were added directly to the normal culturing media for each cell type. After 12-16 hours, media was replaced and cells were incubated for another 24 hours for cells transduced with VSV-pseudotypes, and 48 hours for those transduced with LV pseudotypes. Cells were then fixed with 2% paraformaldehyde (PFA, #43368-9M, Alfa Aesar) and stained with Alexa Fluor 546-conjugated phalloidin (A22283, Invitrogen) and Hoechst (#H3570, Molecular Probes). GFP expression in cells served as a readout for viral transduction.

### Viral titering

To determine functional titers of viral pseudotypes, 3x10^3^ empty BHK21 cells or those expressing dox-inducible Myomaker and/or Myomerger, or EFF-1 (with or without Cre-reporter) were plated in each well of a 96-well tissue culture plate with 500 ng/ml doxycycline. 24 hours later, cells were overlayed with serial dilutions of VSV or LV pseudotypes with 6 μg/ml polybrene and subjected to spinfection by centrifugation at 600xg for 1 hour at 22°C. For VSV pseudotypes, viral titers were determined 24 hours following transduction where cells were fixed with 2% PFA and stained with Hoechst, followed by quantification of GFP^+^ cells using the Cytation™ 5 (BioTek, Winooski, VT, USA). For lentiviral pseudotypes, media was changed 16-18 hours following transduction and cells were analyzed 24 hours later. Cells were fixed with 2% PFA and stained and imaged or trypsinized and GFP^+^ cells were counted using the Countess 3 Automated Cell Counter (Invitrogen, Waltham, MA, USA). Viral titers were then determined using the following formula TU/mL = (Number of cells transduced x Percent fluorescent x Dilution Factor)/(Transduction Volume (mL)).

### Preparation and labeling of extracellular vesicles

Culture media of 10T½ fibroblasts overexpressing Myomaker and/or Myomerger or empty plasmid was collected following two days in exosome-depleted serum and sequentially centrifuged at 1000xg for 10 minutes then 10,000xg for 30 minutes. The supernatants were collected and passed through a 0.22-mm filter (Corning, Corning, NY, USA), followed by ultracentrifugation at 100,000xg at 4°C overnight. Extracellular vesicle pellets were washed with cold PBS and recovered by ultracentrifugation at 100,000xg for 3 hours. For protein detection on exosomes, pellets were lysed in NP-40 lysis buffer and further processed for immunoblotting as mentioned above.

### Functional assessment of Myomaker and Myomerger coated extracellular vesicles

Extracellular vesicle (EV) pellets were resuspended in PBS following final ultracentrifugation and equal concentrations of extracellular vesicle were labeled with lipophilic membrane stain (1,1’-Dioctadecyl-3,3,3’,3’-Tetramethylindocarbocyanine Perchlorate (’DiI’; DiIC18(3)). EVs were washed one time with PBS to remove unincorporated DiI. 5 μg of DiI-labeled EVs were overlayed onto primary myotubes in 8-well chamber slides in corresponding culturing media for 12 hours. Cells were then washed twice with DMEM and replenished with fresh culturing media. 12 hours later, cells were fixed with 2% PFA and analyzed for DiI fluorescence as a readout of extracellular vesicle uptake and also stained with Hoechst and wheat germ agglutinin.

### In vivo delivery of lentiviral pseudotypes

Bald, VSV-G, or Myomaker + Myomerger-pseudotyped lentivirus coding for Luciferase, Cre, or μDystrophin was prepared as described above. For intramuscular injections, 40-80 mL of viral supernatant (per muscle) was first centrifuged for 5 minutes at 2,500 RPM to clear cellular debris and passed through a 0.45 μm SFCA filter. Supernatants (containing the virions) were then concentrated by centrifugation at 10,000 RPM at 4°C for at least 4 hours. The viral pellet was washed once with PBS followed by centrifugation at 10,000 RPM at 4°C for at least 4 hours. The viral pellet was finally resuspended in 25-40 μl PBS and injected into the tibialis anterior or plantaris muscle of the indicated mice using an insulin syringe equipped with a 28-gauge needle. For retro-orbital injections 50-80 ml of viral supernatant (per mouse) was concentrated as mentioned above. The viral pellet was finally resuspended in 60-100 μl PBS and injected retro-orbitally by the Cincinnati Children’s Hospital Medical Center Comprehensive Mouse and Cancer Core.

Some cohorts of mice were subjected to cardiotoxin-induced muscle injury or muscle overload prior to delivery of pseudotyped lentivirus. 35 μl of 10 μM stock of cardiotoxin (EMD Millipore Cat. #217503) was injected into the tibialis anterior muscle and viral injections were administered to 4 days later. Overload of the plantaris muscle was achieved through bilateral synergist ablation of the soleus and gastrocnemius muscles of 3-month-old mice. Briefly, the gastrocnemius muscle was exposed by making an incision on the posterior-lateral aspect of the lower limb. The distal and proximal tendons of the soleus, lateral and medial gastrocnemius were subsequently cut and carefully excised. Viral injections were administered to plantaris muscle 7-days following surgery.

### Immunoblotting

Cells were pelleted at 14,000 RPM for 2 minutes at 4°C and washed with PBS. Viral particles were pelleted from 1 mL of viral supernatant by centrifugation at 10,000 RPM at 4°C for at least 4 hours. Cell or viral pellets were lysed in NP-40 lysis buffer (50 mM Tris-Cl (pH 8.0), 200 mM NaCl, 50 mM NaF, 1 mM dithiotheritol, 1 mM sodium orthovanadate, 1 mM Benzamidine, 0.3% NP-40 and protease inhibitors). 100 μg of protein from cells or total amount of pelleted viral protein from 1 mL of viral supernatant was resolved on 10-15% SDS–polyacrylamide gel electrophoresis and electro transferred onto a nitrocellulose membrane, probed using antibodies against Myomaker (1:500, provided from Dr. Leonid Chernomordik laboratory), Myomerger (1:1000, #AF4580, R&D ), VSV-G (1:1000, #8G5F11, Kerafast), V5 (1:1000, #R96025, Invitrogen), HIV-Gag (p55, p24, p17; 1:1000, #ab63917, Abcam) and GAPDH (1:2000, #MAB374, Millipore) followed by secondary antibodies against rabbit, sheep, or mouse conjugated to Alexa-Fluor 680 (Invitrogen) , DyLight 680, or DyLight 800 (Cell Signaling Technology. Bands were visualized using the Odyssey infrared detection system (LI-COR Biosciences).

### Transmission Electron Microscopy (TEM)

Lentiviral particles were placed on a Formvar Carbon Film (400 mesh, copper, Electron Microscopy Sciences) and processed for electron microscopy by staining with 1% ammonium molybdate. Viral particles were examined under an EM10 C2 microscope (Zeiss, Germany) at 80 kV.

### Muscle histology

For tdTomato analysis, skeletal and non-skeletal muscle tissues were isolated and fixed in 2% PFA/PBS overnight at 4°C then placed in 20% sucrose/PBS at 4°C for at least 4 hours. Tissues were embedded in optimal cutting temperature (OCT) compound (Sakura Finetek, Torrance, CA, USA) and frozen, then 10 μm sections were collected. The sections were mounted with VectaShield containing DAPI (Vector Laboratories, Burlingame, CA, USA).

For analysis of uDystrophin expression, skeletal and non-skeletal myogenic organs were coated with talcum powder and snap frozen in liquid nitrogen. Tissues were then embedded in OCT compound and 10-μm sections were collected. Sections were fixed in ice-cold acetone for 10 minutes, blocked with 2% BSA/PBS for 30 minutes and incubated with: a) anti-Dystrophin (Dy10/12B2) (1:50, #GTX01869, GeneTex) followed by secondary antibody Alexa Fluor 546 (#A-11018, Invitrogen) (Fig. 6E) or b) anti-Dystrophin (1:50, MANEX1011B(1C7), DSHB) conjugated to Alexa Fluor 488 (A20181, Thermo Scientific) (Fig. 6B). Along with dystrophin antibodies, some sections were incubated with anti-Laminin antibodies (#L9393, Sigma Aldrich). All antibodies were diluted in blocking solution and incubated on sections overnight at 4°C. All sections were mounted with VectaShield containing DAPI. All immunostaining in vitro and in vivo was visualized with a Nikon Eclipse Ti inverted microscope with A1R confocal running NIS Elements (Nikon, Tokyo, Japan), and images were analyzed with Fiji (ImageJ, National Institutes of Health, Bethesda, MD, USA).

Masson’s Trichrome staining was performed for analysis of fibrosis in diaphragm muscles using a commercially available kit. 10 μm sections were prepared from frozen diaphragm muscles and were stained following a protocol suggested by the manufacturer (Richard-Allan Scientific).

### Bioluminescence imaging

Mice were injected with 150 mg/kg of D-luciferin (Promega, Cat. # E1605 or Goldbio, Cat. # LUCK-2G) and anesthetized with isoflurane before imaging. The bioluminescence images were acquired 7 min after D-luciferin injection with a rate of one image per 1 minute for 30 minutes using the IVIS spectrum CT in vivo imaging system (PerkinElmer, Cat#128201). Total radiance was measured from the same size of the region of interest (ROI) using Living Image 4.7.3 software (PerkinElmer). The highest captured radiance over imaging time was determined as the peak of the kinetic curve and chosen for analysis.

### FACS analysis

For analysis of pseudotyped lentiviral transduction in mononuclear myogenic cells in vivo, tibialis anterior muscles were harvested and subjected to enzymatic digestion by collagenase and pronase. Mononuclear cell suspensions were incubated with antibodies against α7-integrin (MBL International, Woburn, MA, USA) and Alexa Fluor 488 for detection of myogenic cells and CD45, CD31, Sca1 and Ter119 for negative selection (all Alexa Fluor 700 conjugated, eBiosciences). FACS analysis was performed on a FACSCanto (BD Biosciences) or LSRFortessa (BD Biosciences). Contour plots of data were generated with FlowJo v10.8.1 software.

## QUANTIFICATION AND STATISTICAL ANALYSIS

Quantification was combined from at least two independent experiments, each with technical replicates. At least two independent viral or EV preparations were utilized. Sample sizes and replicates for each experiment are noted in the figure legends. Data were processed and analyzed using Microsoft Excel and GraphPad Prism 9 software. In all graphs, error bars indicate the standard deviation (SD). Data were compared between groups using various statistical tests based on number of groups, normality of data, variance of standard deviations, and multiple comparisons. Specific statistical tests are noted in the figure legends. The criterion for statistical significance was *p< 0.05, **p< 0.01, ***p< 0.001, ****p< 0.0001.

**Figure S1.**
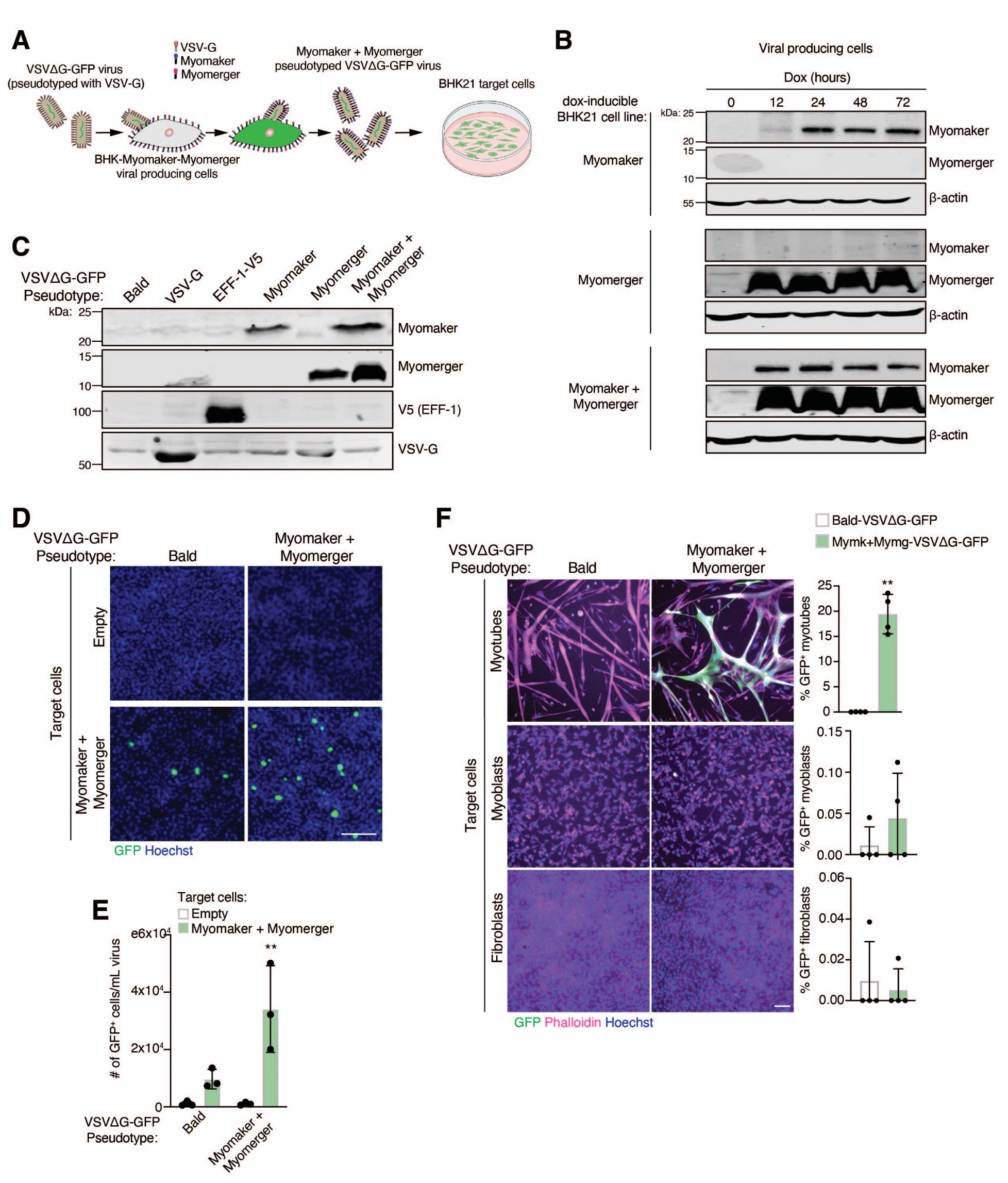
Myomaker and Myomerger on the envelope of VSVs direct fusion toward muscle-fusogen-expressing cells, related to Figure 1. (A) Schematic of the VSVΔG-GFP pseudotyping system. VSVΔG-GFP is used to infect Myomaker^+^ Myomerger^+^ target cells, which will generate a new virus lacking VSV-G but containing Myomaker and Myomerger on the envelope. (B and C) The indicated proteins were detected by immunoblotting from cell lysates (B), or viral supernatants (C). (D) Representative images showing GFP^+^ cells after Bald-VSVΔG-GFP or Mymk+Mymg-VSVΔG-GFP were applied to empty target cells or cells expressing Myomaker and Myomerger. Scale bar, 100 μm. (E) Quantification of transduction units (GFP^+^ cells/mL viral supernatant) from (D). (F) The indicated VSVΔG-GFP pseudotypes were placed on differentiated myotubes, proliferating myoblasts, or fibroblasts and cultures assayed for GFP, and stained for Phalloidin and Hoechst. Representative images are shown. Scale bar, 100 μm. Quantification of the percentage of GFP^+^ cells are shown to the right for each cell type. Data are presented as mean ± standard deviation. Statistical tests used were (E) Two-way ANOVA with Tukey’s correction for multiple comparisons; (F) Unpaired t-test with Welch’s correction; **p<0.01, ****p< 0.0001.

**Figure S2.**
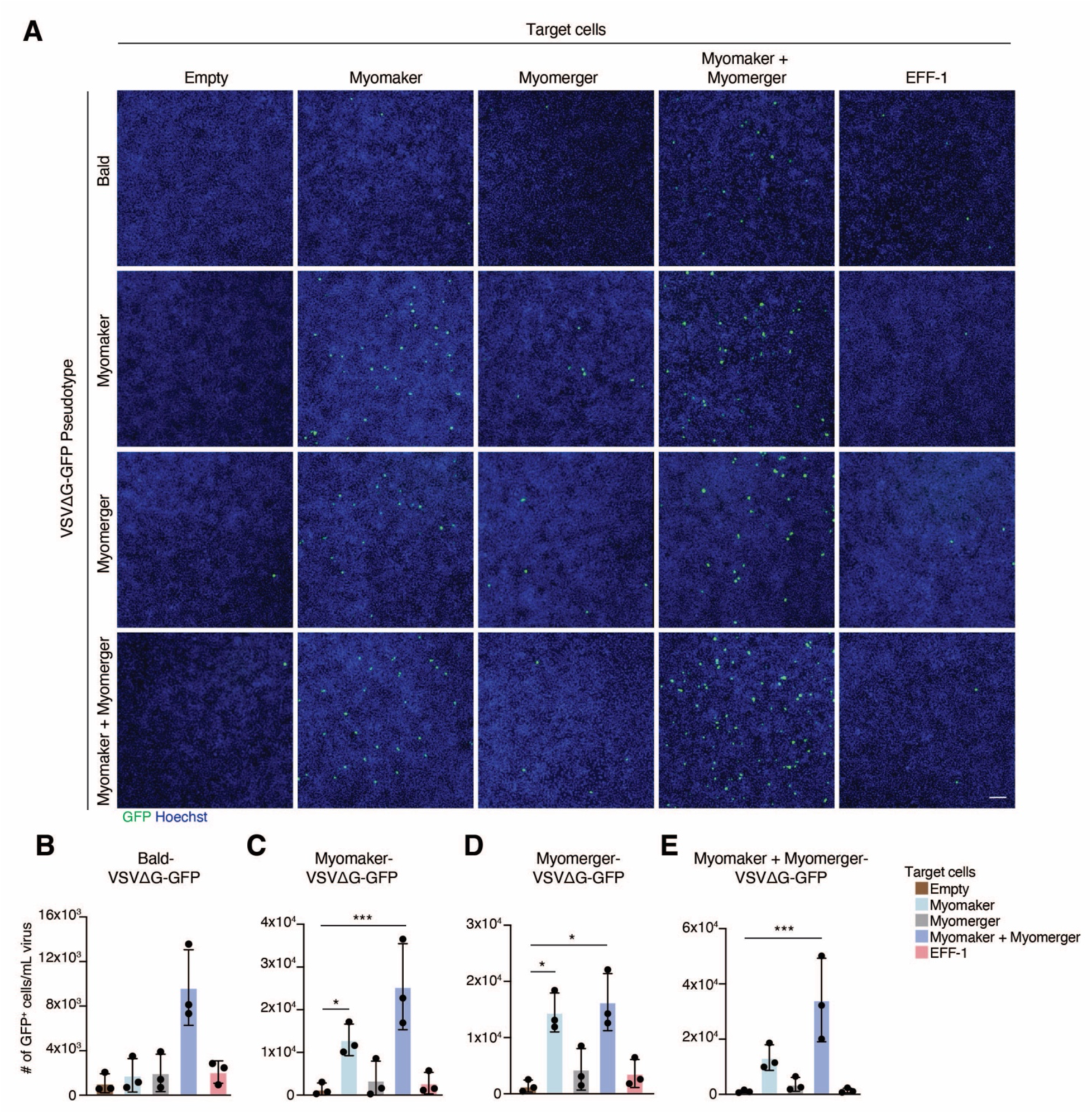
Characterization of the requirement of the fusogens on the viral and cell membranes in the VSVΔG-GFP pseudotyping system, related to Figure 2. (A) VSVΔG-GFP pseudotypes are indicated on the left side and target cells on the top. Representative images showing GFP^+^ cells are shown. Scale bar, 100 μm. (B-E) Quantification of GFP^+^ cells/mL virus as an indicator of transduction efficiencies for each of the pseudotypes on the five different target cells. Note that each graph has different axis magnitude. Data are presented as mean ± standard deviation and combined from three independent VSV preparations. Statistical tests used were (B) repeated measures Friedman test with Dunn’s correction for multiple comparisons; (C-E) repeated measures one-way ANOVA with Tukey’s correction for multiple comparisons; *p<0.05, ***p<0.001.

**Figure S3.**
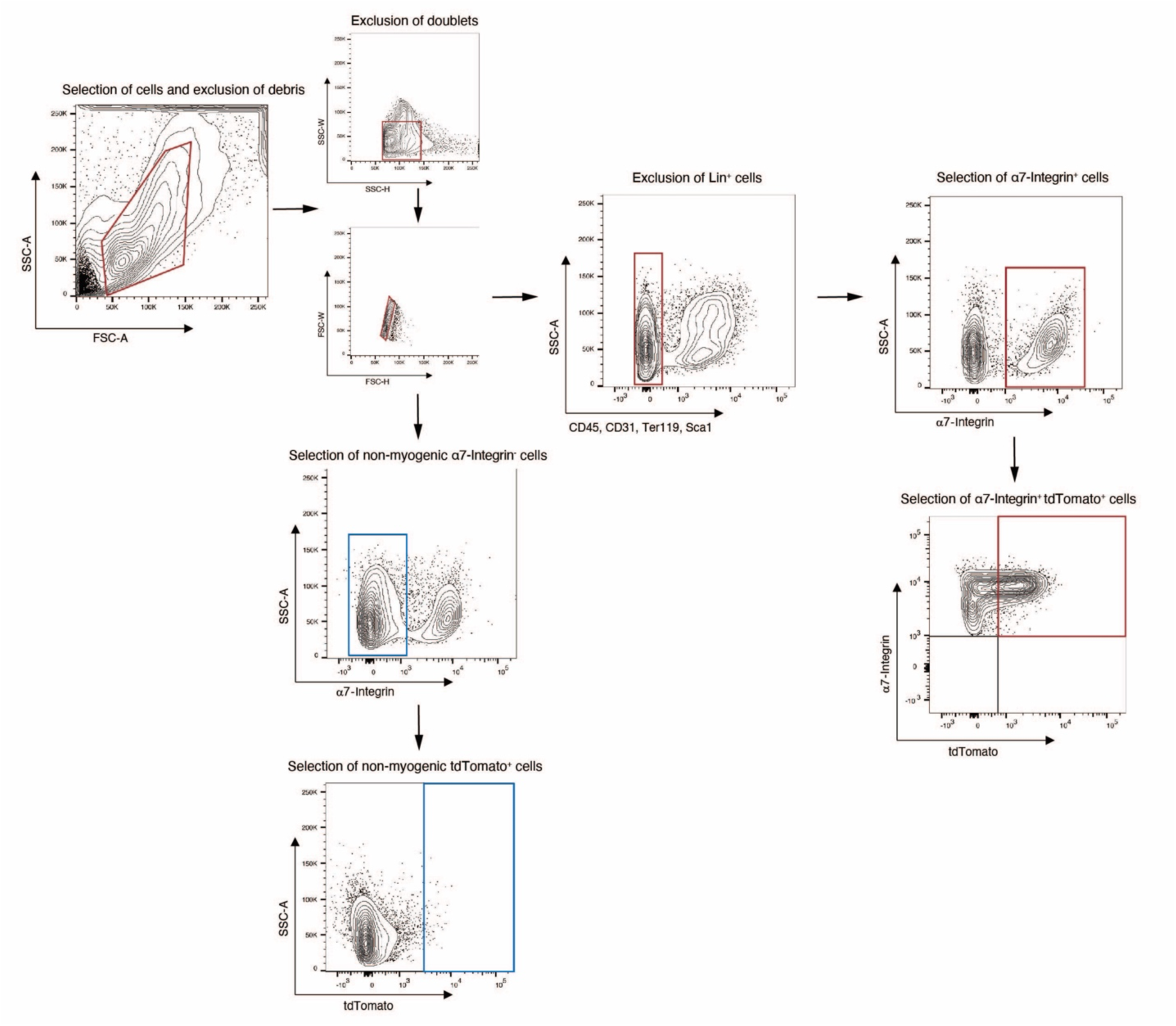
FACS strategy to identify non-myogenic cells and myogenic progenitors containing tdTomato, related to Figure 4. (A) Mononuclear cells were digested from tibialis anterior muscles and stained with antibodies for CD45, CD31, Ter119, Sca1, and α7-Integrin. All plots are representative from two independent experiments each with at least three samples per group.

**Figure S4.**
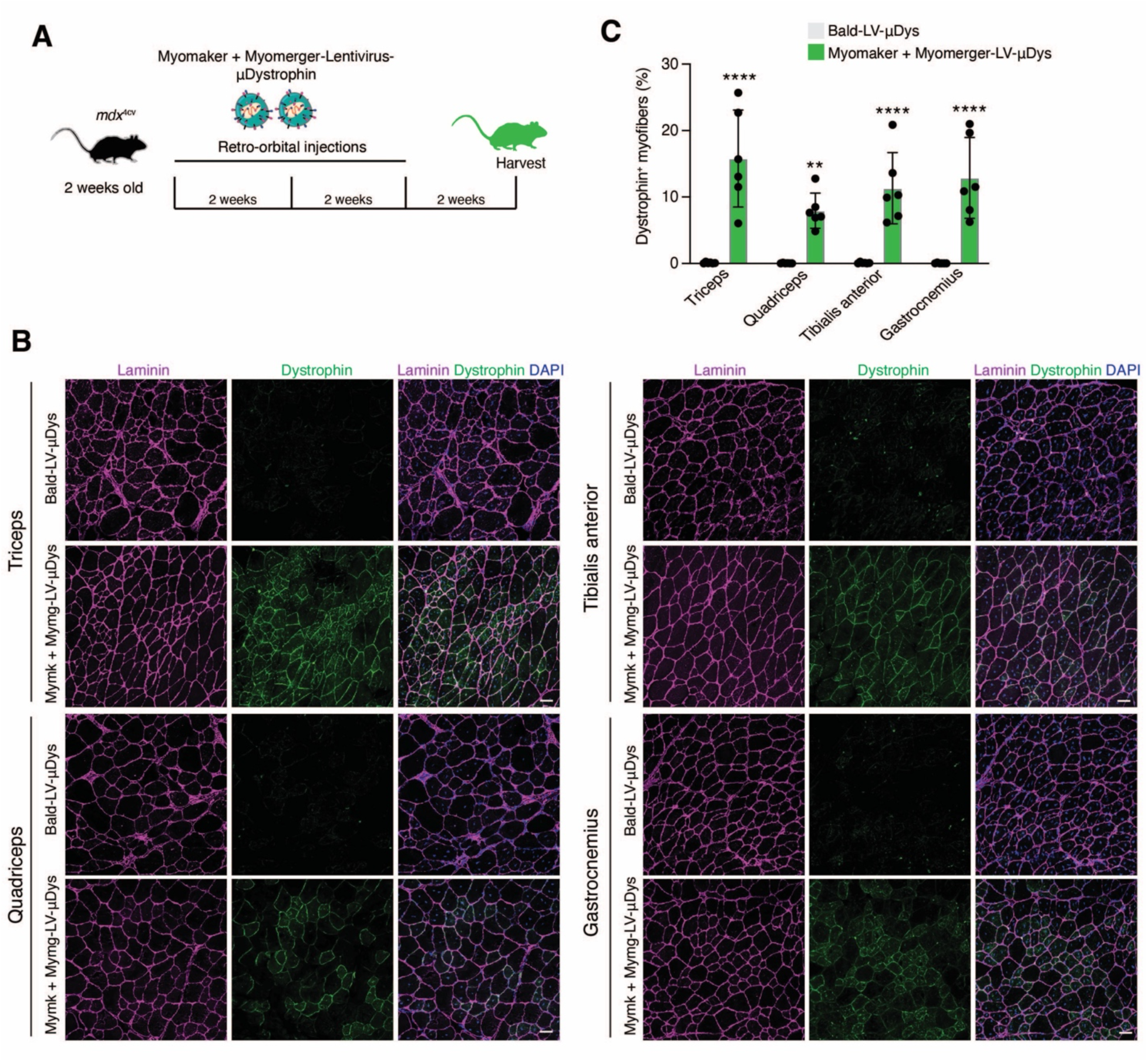
μDys in limb muscles after systemic delivery of Mymk+Mymg-LVs, related to Figure 6. (A) Experimental design for systemic delivery of Bald-LV-μDys or Mymk+Mymg-LV-μDys. (B) Immunostaining for dystrophin in *mdx*^4cv^ limb muscles treated with Bald-LV-μDys or Mymk+Mymg-LV-μDys through retro-orbital injections. Scale bars, 50 μm. (C) Quantification of μDys^+^ myofibers. Data are presented as mean ± standard deviation and combined from two independent LV preparations. Statistical test used was unpaired t-test with Welch’s correction; **p<0.01, ****p<0.0001.

**Figure S5.**
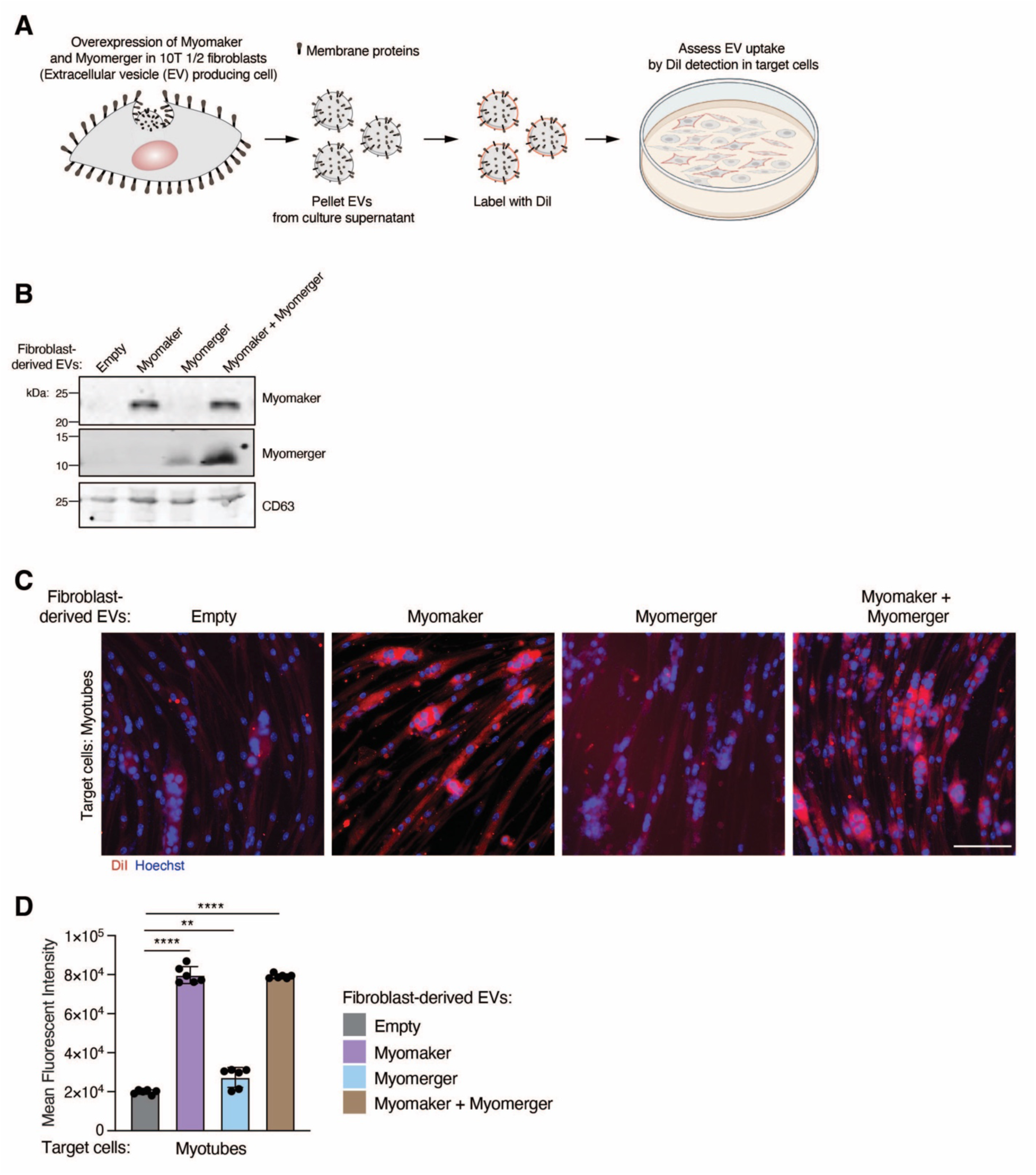
Extracellular vesicles with Myomaker and Myomerger increase delivery to myogenic cells, related to Figure 1. (A) Schematic of experimental design to produce extracellular vesicles (EVs) from fibroblasts transduced with an empty virus, Myomaker, Myomerger, or Myomaker and Myomerger. EVs were harvested by ultracentrifugation, labeled with DiI, and applied to target cells. (B) Western blot from the indicated fibroblast-derived EVs to detect the presence of Myomaker, Myomerger, and CD63 (EV marker). (C) Representative images of myotubes after treatment with various EVs. Cells were imaged for DiI as an indicator of delivery by EVs, and Hoechst (nuclei). Scale bar, 100 μm. (D) Quantification of the mean fluorescent intensity for DiI from (C). Data are presented as mean ± standard deviation and combined from at least three independent extracellular vesicle preparations. Statistical test used was (D) One-way ANOVA with Tukey’s correction for multiple comparisons; **p<0.01, ****p<0.0001.

